# Subtle alteration in transcriptional memory governs the lineage-level cell cycle duration heterogeneities of mammalian cells

**DOI:** 10.1101/2024.05.23.595451

**Authors:** Kajal Charan, Sandip Kar

## Abstract

Cell cycle duration heterogeneities within a malignant tumor significantly reduce the therapeutic efficacy of cancer treatment. However, identifying factors governing such heterogeneities remains challenging. Here, we demonstrate that correlated fluctuations of the inherited transcription rates during cell cycle progression in combination with the random resetting of them during mitosis are sufficient to account for the experimentally observed cell cycle duration correlation patterns for daughter-daughter, mother-daughter, and cousin-cousin cell lineage pairs. The model elucidates that the variations in the transcriptional inheritance pattern dictate the extent of the cousin-mother inequality phenomenon. It further predicts that the longer the cell cycle duration, the lower will be the lineage pair correlations. However, higher S-G2-M phase duration will only increase daughter-daughter and mother-daughter correlations, hence, reducing the extent of cousin-mother inequality. Overall, the model shows how alteration of the inherited transcriptional fluctuations across lineages finetunes the lineage-level heterogeneities.

## Introduction

Cells within a malignant tumor demonstrate a high degree of heterogeneity in their proliferation pattern ^1–5^. This makes sophisticated chemotherapeutics ineffective in eradicating cancerous cells ^6–9^. Mammalian cells, even under culture conditions, display significant variations in their cell cycle period and phase durations ^1,10,11^. Recent single-cell imaging studies ^12–17^ have revealed interesting cell cycle duration-related correlation patterns at the lineage level for various mammalian cell types. The sister pairs (D-D) are found to be highly correlated in their cell cycle duration, and mother-daughter (M-D) and cousin pairs (C-C) exhibit relatively poorer correlations in comparison to the sister pairs. Intriguingly, there are instances where the C-C cell cycle duration correlation is higher compared to M-D pairs (termed cousin-mother inequality, **CMI**) ^12–14^, and sometimes M-D correlation is either higher or comparable to the C-C correlations ^15,16,18^. These observations led to many interesting questions which remain unanswered. For example; (i) Why do we observe such kind of cell cycle duration heterogeneities? (ii) Why are the sister pairs highly correlated? (iii) How C-C correlations can exist when M-D pairs are poorly correlated in the first place? (iv) Why **CMI** does not consistently hold for all cell types? Answering these questions will unravel why such kind of lineage-level correlations are conserved across different mammalian cell types and what causes such subtle variations in these correlation patterns.

In literature, various modeling approaches have been employed to explain the origin of such lineage-level correlation patterns for specific cell types to explain specific experimental findings ^1,13–16,18^. However, The origin of such kind of duration heterogeneities at the lineage level remains vaguely understood as each of these studies has put forward different reasoning to elucidate a universal phenomenon without addressing several other associated intricate features of the cell cycle duration heterogeneities. In a recent study, ^18^ explained the correlation pattern as a universal feature across various cell types. They suggested that some heritable cell cycle regulatory factors play the governing role in generating diverse correlation patterns in lineage pairs. How exactly such factors are inherited and how this kind of inheritance is modulated during the cell cycle to impact the correlation patterns remains unclear.

In this article, we analyze a simple cell cycle model that elucidates all the correlation-related aspects of the cell cycle duration heterogeneities at the lineage level in a comprehensive way. The model hypothesizes that the daughter cells within a lineage inherit the transcription rate of various cell cycle regulatory genes from their mother ^19–22^ and these transcription rates then fluctuate in a correlated manner during cell cycle progression. Our model demonstrates that the variations of noise strengths and correlation times of such kind of correlated transcriptional fluctuations can explain the complex lineage-level cell cycle duration correlation patterns as observed experimentally for various mammalian cell types. The model further delineates how the cell cycle period and the individual phase durations affect the cell cycle duration heterogeneities in the presence of correlated transcriptional fluctuations.

## Model

Our model is an improvised version of a generic mammalian cell cycle model developed by Tyson and Novak ^23^. This model includes three regulatory proteins, CycB, Cdh1, and Cdc20 (**Fig.1A**), whose interactions control the entry into mitosis and subsequent exit from the cell cycle. Specifically, CycB and Cdh1 have an antagonistic relationship that maintains a low CycB level during the early G_1_ phase as the Cdh1 sustains a high level. In the current model, we have assumed that an increase in the externally added growth factor (GF) activates the CycB which can phosphorylate the Cdh1 to make it inactive and cells eventually transit from G_1_ to S-phase^24,25^. However, a higher level of CycB activates the transcription of Cdc20 which dephosphorylates Cdh1 and facilitates the cell cycle exit (Detailed model in **SFig.1**). Here, our endeavor is to follow several lineages of cells (**Fig.1B**) for a specified time period (like 72 hours) computationally as happens during live-cell imaging experiments ^12,13,15^. We have implemented the cell division event when CycB reaches a certain threshold level during late mitosis ^23^ (**SFig.2)**. We have assumed that all the cellular components in the mother cell get equally distributed between the two daughter cells during the division event.

**Fig. 1.**
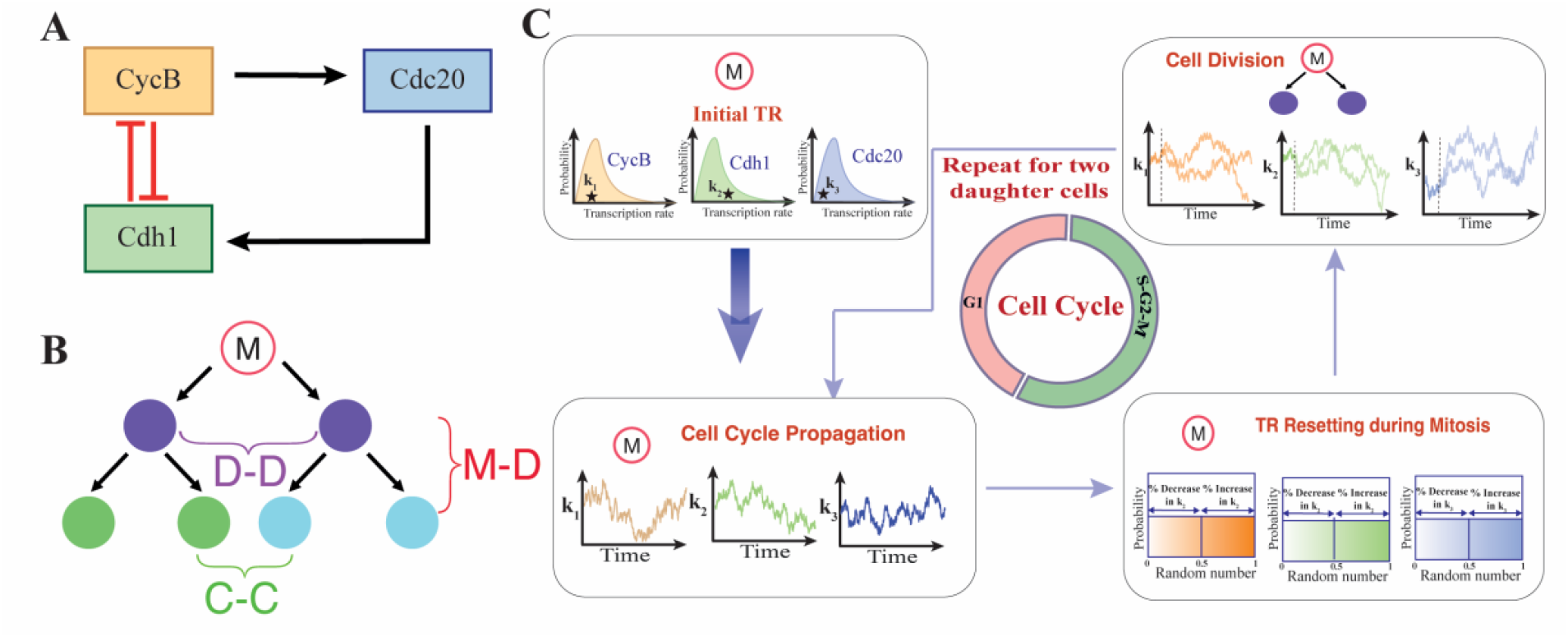
The proposed generic model and the simulation protocol to capture the lineage level cell cycle duration variabilities. (A) The proposed cell cycle regulatory network ^23^. (B) A representative cell lineage tree. (C) The schematic numerical protocol for simulating cell cycle lineages by introducing transcriptional fluctuations in cell lineages and during the cell cycle progression.

Importantly, we have included the dynamics of mRNAs for each regulatory protein in the model proposed by Tyson et al. to introduce the concept of correlated transcriptional inheritance across cellular generations ^19,20,26–30^. In this regard, a few experimental observations are extremely critical. For example; Phillips et al. have observed that despite significant gene-level fluctuations, the mean transcriptional activity is restored in daughter cells from mothers within lineages in mouse embryonic stem cells ^20^. Rosenfeld et al. have demonstrated that slow fluctuations in protein production rate have a correlation timescale equivalent to the duration of a cell cycle ^26^. While Sigal et al. have shown that in mammalian cells these correlation timescales in protein production can be even more than one cell cycle duration and vary in a gene-dependent manner ^19^. Considering these observations, in our model (**Table 1-3**) we have assumed that daughter cells inherit the transcription rates of all genes from their mother. However, these transcription rates have certain correlated fluctuations during the cell cycle progression with non-zero correlation timescales.

The time series of this correlated noise is generated using the Ornstein-Uhlenbeck process as described in **Eq.1**.^30–32^

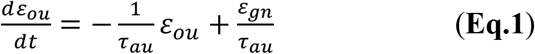

Where, *ε*_*gn*_ is the delta correlated white noise with zero mean, i.e.,

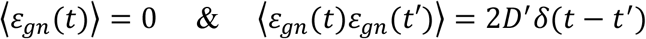

Here, *ε*_*ou*_ is now the exponentially correlated noise which has a positive autocorrelation time *τ*_*au*_ and follows the relations 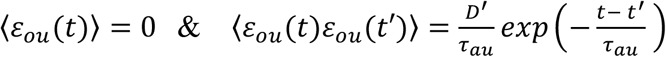. In our simulation, we generated the Gaussian white noise by picking random numbers from the Gaussian distribution which have noise strength *D* = 4*D*^′^Δ*t* (Δ*t* is the time step of simulation) (Fox et al., 1988).

To perform the lineage-level simulations, at *t* = 0, we randomly pick up a mother cell for each lineage. The transcription rate of a specific gene for a representative mother cell is initially chosen from a log-normal distribution with a deterministic mean transcription rate for the same gene and a 5% coefficient of variation (CV)^33,34^ (**Fig.1C**). These transcription rates are then propagated with colored noise having non-zero correlation timescales during cell cycle progression^30,31^. Additionally, we have incorporated the transcription rate variations that happen during the Mitotic phase, resulting from chromatids segregation into the two daughter cells ^21,22,35,36^. Upon cell division, the two daughter cells inherit all transcription rates from their mother ^19^ (for details see **Methods**).

## Results and Discussions

### Alteration in transcriptional memory reconciles diverse cell cycle duration correlation patterns in lineage pairs

Following the proposed simulation scheme (**Fig. 1C**), we have been able to capture the diversity in the cell cycle duration heterogeneities across cell lineages (**Fig.2**). First, we demonstrate how transcriptional memory can generate correlation among lineage pairs. Our simulation at a fixed noise strength (*D* = 0.03) with varying autocorrelation time (*τ*_*au*_) indicates that as *τ*_*au*_ increases, there is a rise in correlation among all lineage pairs (more in M-D and D-D pairs, less in the C-C pair) (**Fig.2A**) which can be attributed to the higher transcriptional memory. Interestingly, with increasing *τ*_*au*_, an intriguing pattern emerges. At lower values of *τ*_*au*_, the M-D correlation is less than the C-C correlation, however, beyond a threshold value of *τ*_*au*_, the M-D correlation supersedes C-C correlation (**Fig. 2A**). This suggests that enhancement in transcriptional memory can diminish the initially observed **CMI** by having a greater impact on the correlation between M-D pairs than the correlation between C-C pairs. Experimentally it has been exhibited that ^19^ the autocorrelation time and the extent of variability in transcription rates vary notably across various genes. Thus, we simulate a similar simulation by varying the noise strength (*D*) while keeping the *τ*_*au*_ fixed at 10 hours. The simulations reveal that with increasing *D*, all the lineage pair correlations decrease, and beyond a specific *D* value, there is an emergence of **CMI** (**Fig.2B)**. It is important to emphasize that we have performed these calculations assuming a 24-hour cell cycle period (with 10 hours G_1_ and 14 hours S-G_2_-M durations).

**Fig. 2.**
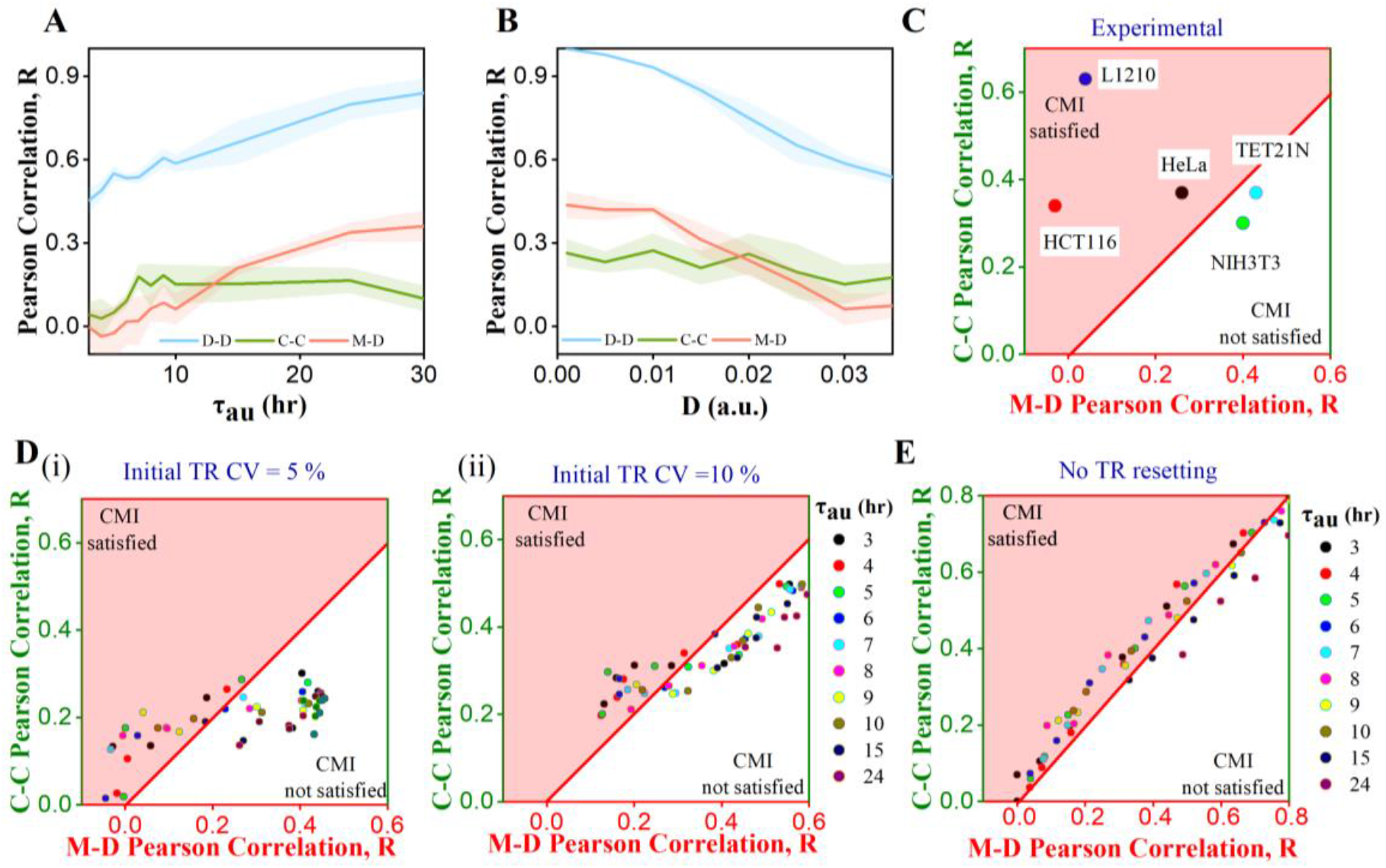
Different Lineage correlation patterns emerge due to variabilities in transcriptional memory. Correlation in D-D, M-D, and C-C with change in **(A)** *τ*_*au*_ = 3 hours to 30 hours (fixed *D* = 0.03) and **(B)** at varying *D* (fixed *τ*_*au*_ =10 hours). Plot of M-D cell cycle duration correlation against C-C correlation; **(C)** from experimental literature for different cell lines.^12–16^, **(D)** from our simulations for *τ*_*au*_ ranging from 3 hours to 30 hours at different *D* for initial Transcription rate (TR) distributions with CV (i) 5% and (ii) 10%, and **(E)** from our simulations without TR resetting at mitosis. The shaded area shows the region where **CMI** is satisfied. All simulations are performed for 3 replicates each for 300 cells. Error bar cloud in **Fig 2A & 2B** represents the S.E.M. of 3 replicates.

To comprehensively understand the collective influence of *τ*_*au*_ and *D* on the variabilities in lineage pair correlations, we have performed simulations across a spectrum of *τ*_*au*_ and *D* values (**Fig.2D(i)**). Our simulations qualitatively reconcile the experimentally observed correlation pattern among M-D and C-C lineage pairs for different cell types (**Fig.2C**) ^12,14,16–18^. **Fig.2D(i)** further delineates that for certain combinations of *τ*_*au*_ and *D*, we will observe **CMI** as seen in experimental studies ^12,14,18^. Further, it has been observed that the CV for the initial log-normal distributions for various transcription rates from which the initial set of transcription rates are randomly selected for the starting mother cells in each lineage plays a substantial role in determining the correlations of cell cycle duration in D-D, M-D and C-C pairs ^20^.

Intriguingly, our simulations predict that increasing the CV values for all the log-normal distributions for the transcription rates leads to greater Pearson’s correlation in lineage pairs (**Fig.2D(ii)**) which is purely statistical in nature. This increase can be attributed to the greater linear rise of the covariance term over the increase in variances of cell cycle durations. This suggests that for cancerous cells with a greater possibility of transcription rate variability across cellular populations, we will observe more correlated cell cycle durations in the lineage pairs.

At this point, we further analyze the effect of the additional sources of variability alongside the correlated transcriptional fluctuations. In our algorithm, we have considered the transcription rate variations during mitosis, which quantitatively accounts for the epigenetic modifications occurring during sister chromatid separation among the two daughter cells ^21,22,35,36^. Interestingly, we found that the exclusion of this effect in our simulation does not affect the D-D correlation, however, it increases the M-D and C-C correlations significantly, which is not observed in the experiments (**Fig.2E**). This foretells that such kind of transcriptional variation during mitosis plays an essential role in reducing the M-D and C-C correlations without influencing the high D-D correlations. Thus, our model simulations provide qualitative insight into how the finetuning of the transcriptional memory across lineage and during the cell cycle generates diverse lineage-level correlation patterns.

### Lineage-level correlation decreases with increasing cell cycle duration leading to an extended region of CMI

In **Fig.2**, simulations are performed assuming a ∼24 hours cell cycle duration. However, the cell cycle duration in mammalian cells can vary from ∼16 to 40 hours, which can lead to distinct correlation signatures. Herein, we investigate how alterations in cell cycle duration can influence these correlation patterns under specific transcriptional memory-related parameters. The simulation results indicate that correlation decreases among all lineage pairs as the cell cycle duration is increased monotonically from 16 hours to 28 hours maintaining a proportional increase in G_1_ and S-G_2_-M phase durations (**Fig.3A**).

**Fig. 3.**
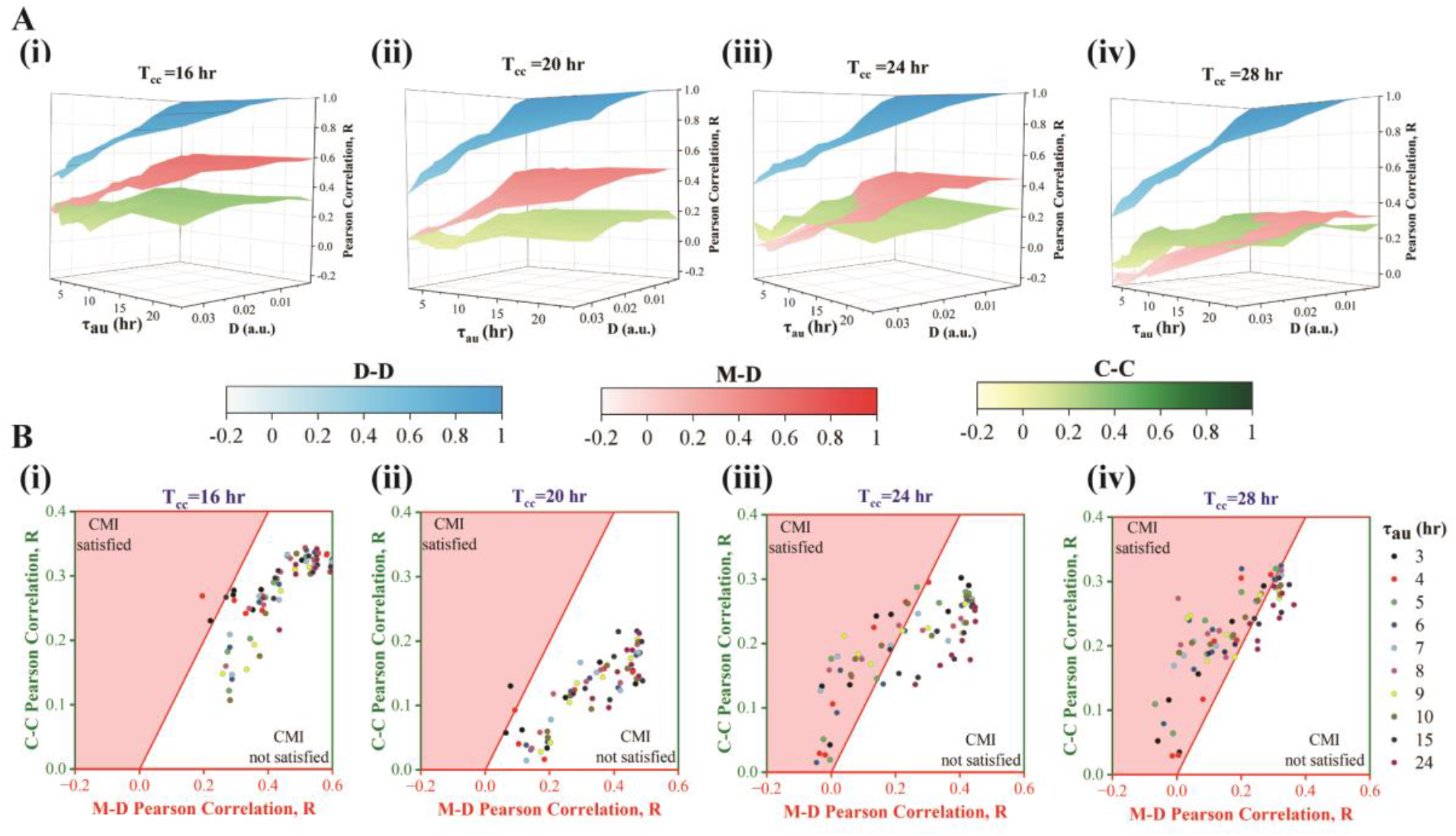
Lineage pair correlation patterns for varied cell cycle durations. **(A)** Surface plots of cell cycle duration correlation patterns (D-D, M-D, and C-C) as a function of ***τ***_***au***_ and ***D*** for **(i)16** hour, **(ii) 20** hour, **(iii) 24** hour, and **(iv) 28** hour cell cycle period. **(B)** M-D Vs C-C correlation plot as a function of ***τ***_***au***_ and ***D*** for **(i)16** hour, **(ii) 20** hour, **(iii) 24** hour, and **(iv) 28** hour cell cycle period.

This drop in correlation occurs because the effect of transcriptional memory diminishes with increasing cell cycle durations. These findings qualitatively corroborate the experimental study by Kuchen et al. ^15^ where they prolonged the cell cycle duration by suppressing Myc expression in neuroblastoma cells, which resulted in a decrease in the correlation of lineage pairs. Intriguingly, **Fig.3** depicts that for cells having faster cell cycle period (16 hours or 20 hours, **Fig. 3A(i-ii)**), M-D correlation consistently remains higher than C-C correlation, and **CMI** (**Fig. 3B(i-ii))** is almost non-existent. However, for cell types having slower cell cycle durations (24 hours or 28 hours, **Fig. 3A(iii-iv)**), **CMI** appears (**Fig. 3B(iii-iv)**) for a wider range of *D* and *τ*_*au*_. Overall, our simulations foretell that in cell types with higher cell cycle durations, all the lineage correlations will decrease, however, the relative decrease in M-D correlation will be higher which causes the distinct appearance of **CMI**.

### Modulation in G_1_ and S-G_2_-M phase durations alters lineage-level correlation patterns

In the previous section, we explored how variations in cell cycle duration impact heterogeneity among cell lineage pairs by proportionately adjusting the G_1_ and S-G_2_-M phase durations to vary the cell cycle period. How does the alteration in G_1_ and S-G_2_-M phase durations for a fixed cell cycle period influence the lineage-level correlation patterns? We investigated this question by performing a simulation for two representative cell cycle periods (16 hours and 28 hours) with two different combinations of phase durations (low G_1_ and high S-G_2_-M, and high G_1_ and low S-G_2_-M). Our findings indicate that an extended S-G_2_-M phase (independent of the cell cycle period) increases the M-D correlation (**Fig.4A** and **Fig.4C**) due to mothers spending more time with the same set of transcription rates that are subsequently inherited by daughter cells following transcription rate variations at late M-phase. In contrast, cousin correlation remained unaffected by the variation in phase durations for faster cell cycle period (**Fig.4A**), however, with increasing S-G_2_-M duration for the longer cell cycle period, the C-C correlation seems to be increasing (**Fig.4C**). Importantly, the extent of **CMI** decreases (**Fig. 4A**) with increasing S-G_2_-M phase duration for 16-hour cell cycle duration but it reduces marginally (**Fig.4C**) for 28-hour cell cycle period.

**Fig. 4.**
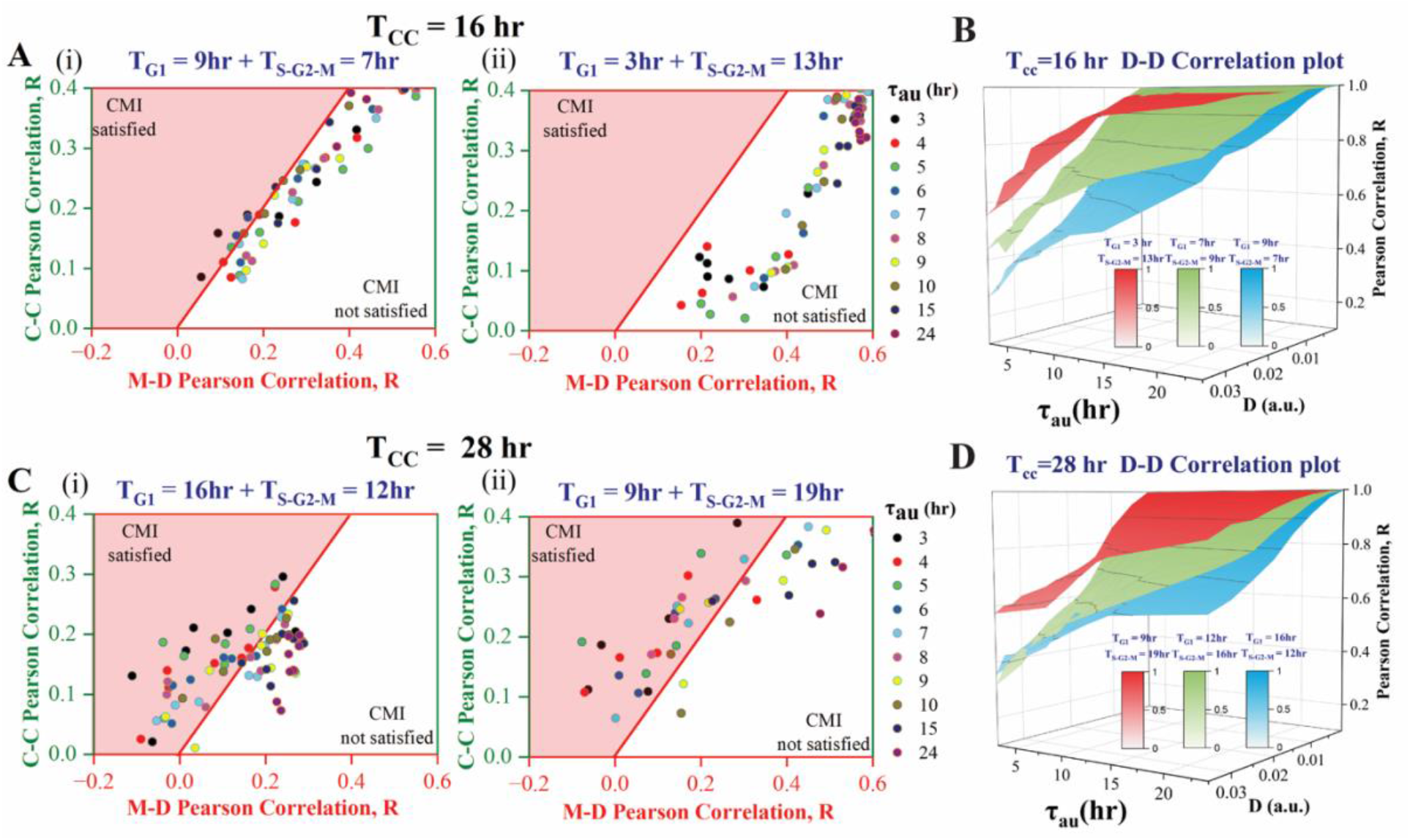
Change in the relative proportion of G_1_ and S-G_2_-M phase durations alters the correlation patterns. **(A)** The plot of M-D and C-C correlation for a cell cycle duration of 16 hours with G_1_ and S-G_2_-M durations **(i)** 9 hours and 7 hours, and **(ii)** 3 hours and 13 hours, respectively. **(B)** D-D correlation plots with varying*τ*_*au*_ and *D* for different proportions of G_1_ and S-G_2_-M phases for a 16-hour cell cycle period. **(C)** Plot of M-D and C-C correlation for a cell cycle duration of 28 hours with G_1_ and S-G_2_-M durations **(i)** 16 hours and 12 hours, and **(ii)** 9 hours and 19 hours, respectively. **(D)** D-D correlation plots with varying*τ*_*au*_ and *D* for different proportions of G_1_ and S-G_2_-M phases for the 28-hour cell cycle period.

Further, we have observed that independent of the cell cycle period, shorter S-G_2_-M duration produces a phase plane distribution having a condensed set of C-C and M-D correlation points (**Fig. 4A(i)** and **Fig. 4B(i)**) as the correlations in lineage pairs are being less affected by the variations in *τ*_*au*_ and *D*. As the S-G_2_-M duration increases, the C-C and M-D correlation points in the phase plane become more dispersed (**Fig. 4A(ii)** and **Fig. 4B(ii)**). Accordingly, our model simulations predict that the longer S-G_2_-M phase duration causes greater M-D correlation and destroys the **CMI** (**Fig.4A** and **Fig.4C**).

Interestingly, the D-D correlation increases with increasing S-G_2_-M phase duration independent of the overall cell cycle period (**Fig.4B** and **Fig.D**). Our simulations reveal that higher S-G_2_-M phase duration (with a fixed cell cycle duration) is indeed lowering the overall variabilities in cell cycle durations at the population level (**SFig.3**) which leads to the enhancement in the D-D and M-D correlations and reduces the **CMI** (**Fig.4**). These model predictions can be verified experimentally by altering specific phase durations implementing various stress conditions ^37^.

## Discussion and Conclusion

Identifying the factors regulating the cell cycle duration heterogeneities in mammalian cells is an important and challenging task. In this regard, recent studies at the single-cell level have provided varied explanations for the lineage-level heterogeneities ^12–18^. Here, we have modified and analyzed a simple generic cell cycle network ^23^ model to understand the origin of lineage-level cell cycle duration heterogeneities in mammalian cells.

Employing our modeling analysis, we have been able to address some of the important open-ended questions in the field of cell cycle duration heterogeneity at the lineage level. First, our model unravels that the correlated variation of the inherited transcriptional rates across generations ^19,20,26^ and its modulation during cell cycle progression are sufficient to qualitatively capture most of the experimentally observed ^16,17,19,20,26^ lineage level correlation patterns (**Fig.2**) in cell cycle durations without taking any other biological factors into consideration. Intriguingly, the model predicts that cell types having higher transcriptional variability across generations (for example cancerous cell lines or stem cells that are prone to differentiate) will have a greater probability of showing stronger cell cycle duration correlations in lineage pairs (**Fig.2D**). Thus, a correlated variation of the inherited transcriptional rates makes the cell cycle duration in daughter pairs highly correlated (**Fig.2A-B**), however, the modification of the transcription rates during mitosis ^35,38^ makes M-D and C-C pairs less correlated than D-D pairs (**Fig.2D-E**). Importantly, depending on the nature and extent of the correlated fluctuations of the inherited transcription rates, the cousin-mother inequality as observed in experiments ^12–16,18^ can be realized in our simulations (**Fig.2A-B**). This suggests that C-C cell cycle duration correlation can still exist due to subtle adjustments in the correlated transcriptional fluctuations during cell cycle progression even if the M-D correlation is low.

Our model further substantiates that the variation in the correlated transcriptional fluctuations finetunes the extent of cousin-mother inequality for different mammalian cells ^12,13,15^ with varied cell cycle duration (**Fig.3**). It demonstrates that cell types with higher cell cycle periods will show a reduced level of correlation in all lineage pairs (**Fig.4**) ^15^. This is because the inherited transcriptional memory will diminish to a greater extent in these cells due to longer cell cycle duration, however, will extend the region of **CMI**. Finally, our model analysis elucidates that a higher extent of S-G_2_-M phase duration will not only increase the D-D correlations, it will also lower the extent of **CMI** by increasing the M-D correlations (**Fig.4**). These model predictions can be easily tested via experiments as these results seem in line with a recent study ^37^ where it has been demonstrated that the presence of heritable and uncoupled factors that maintain a high correlation among siblings while keeping individual phases of the cell cycle uncoupled.

Overall, our modeling study exhibits that the inheritance of the transcription rates across cell lineages and their correlated fluctuations during cell cycle progression can explain the diverse lineage-level correlation patterns in mammalian cells. The model due to its simple nature fails to quantitatively obtain the exact values of the correlations in a cell-type specific manner. However, it provides a qualitative explanation under varied cell cycle periods and phase durations which can be tested experimentally. We believe that these insights will be useful in modifying the cell cycle duration heterogeneities and finding therapeutic relevance.

## Materials and methods

### Simulation in cell lineages

The differential equations, parameter values, and initial conditions given in Table 1-3, respectively, are used for simulation. These differential equations are solved in MATLAB ode solver (ode15s). Further to follow cell lineages, we adapted the methodology similar to Govindaraj et al., where we performed simulations for 72 hours up to three generations ^12^. At cell division (at some specific value of CycB), all variables were divided equally, and the simulation continued for two cells with similar values ^23^.

Cell cycle duration is calculated from the cell’s birth to its division time. To calculate the G1 phase duration, we added an additional variable 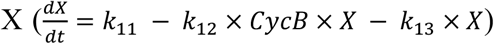. In our simulation, to calculate the precise G1-S transition, we considered X degradation by CycB. The time difference between X peak and cell birth is G1 duration.

All simulations were performed for 100 lineages, and the cells finishing their cell cycle before simulation time were considered. All pairs of Sisters, Mother-daughter, and cousins were separated and used to calculate correlations from the initial 300 cells of each simulation.

### Significance Analysis

We calculated the p-value after performing Fisher’s z transformation on Pearson coefficient (R) values to see if two correlation coefficients differed significantly from one another. From 500 bootstrap random samples, we determined the R-value for each cell subpopulation in a replicate. (Supplementary file)

## Acknowledgments

Thanks are due to UGC for providing the UGC-CSIR-JRF (NTA Ref. No: 191620004555) fellowship to KC. This work is supported by the funding agency **SERB, India** (Grant no. **CRG/2023/002165**). We thank Dr. Amitava Giri and Dr. G. Vinodhini for their suggestions during various stages of this study.

