## supplementary file for "Subtle alteration in transcriptional memory governs the lineage-level cell cycle duration heterogeneities of mammalian cells"

**Table 1 - Ordinary Differential equations governing the cell cycle regulatory network**

| $\frac{dCycBm}{dt}=$ | $\frac{\left( k_{1m}+\varepsilon_{ou1} \right)\times GF}{K_{mm}+k_{eff}*GF}-k_{1dm}\times CycBm$ | **1** |
| --- | --- | --- |
| $\frac{dCycB}{dt}=$ | $k_{1}\times CycBm-k_{2a}\times CycB-k_{2b}\times CycB\times Cdh1$ | **2** |
| $\frac{dCdc20m}{dt}=$ | $\left( k_{5am}+\varepsilon_{ou2} \right)+\frac{\left( k_{5bm}\boldsymbol{+}\varepsilon_{ou3} \right)}{k_{5cm}+GF \times J_{5c}}\times\frac{CycB^{n}}{J_{5}^{n}+CycB^{n}}-k_{5dm}\times Cdc20m$ | **3** |
| $\frac{dCdc20T}{dt}=$ | $k_{5a}\times Cdc20m-k_{6}\times Cdc20T$ | **4** |
| $\frac{dCdc20A}{dt}=$ | $k_{7}\times IEP\times\frac{Cdc20T-Cdc20A}{J_{7}+Cdc20T - Cdc20A}-k_{8}\times Mad\times\frac{Cdc20A}{J_{8}+Cdc20A} -k_{6}\times Cdc20A$ | **5** |
| $\frac{dCdhm}{dt}=$ | $\left( k_{3m}+\varepsilon_{ou4} \right)-k_{3dm}\times Cdhm$ | **6** |
| $\frac{dCdht}{dt}=$ | $k_{3a}\times Cdhm-k_{3dt}\times Cdht$ | **7** |
| $\frac{dCdhA}{dt}=$ | $\left( k_{3}+k_{3b}\times Cdc20A \right)\times\frac{Cdht-Cdh1}{J_{3}+Cdht-Cdh1}-k_{4}\times CycB\times\frac{Cdh1}{J_{4}+Cdh1}-k_{3dt}\times Cdh1$ | **8** |
| $\frac{dIEP}{dt}=$ | $k_{9}\times CycB\times(1-IEP)-k_{10}\times IEP$ | **9** |

**Table 2 -** **Parameter values used for Simulations**

| **Abbreviated Form** | **Full Description** | **Parameter value** | **Unit**  **(c=concentration**  **t=hour)** |
| --- | --- | --- | --- |
| $k_{1m}$ | Transcription rate of CycB mRNA | $0.0037\times d$ | c t^-1^ |
| $k_{1dm}$ | Degradation rate of CycB mRNA | $0.058 \times d$ | t^-1^ |
| $k_{1}$ | Synthesis rate of CycB | 0.4 $\times d$ | t^-1^ |
| $k_{2a}$ | Degradation rate of CycB | 0.04 $\times d$ | t^-1^ |
| $k_{2b}$ | The cdh1-dependent degradation rate of CycB | 2.0$\times d$ | (c t)^-1^ |
| $k_{3a}$ | The synthesis rate of Cdh1 total | 1.0 $\times d$ | t^-1^ |
| $k_{3b}$ | Cdc20 dependent activation rate of Cdh1 | 4.3 $\times d$ | (c t)^-1^ |
| $k_{4}$ | CycB induced degradation rate of Cdh1 active form | 40.0 $\times d$ | t^-1^ |
| $j_{3}$ | Michalis Menten coefficient for Cdc20 dependent activation rate of Cdh1 | 0.04 | c |
| $j_{4}$ | Michalis Menten coefficient for CycB induced degradation rate of Cdh1 active form | 0.04 | c |
| $k_{3m}$ | Transcription rate of Cdh1 mRNA | 0.5 $\times d$ | c t^-1^ |
| $k_{3dm}$ | Degradation rate of Cdh1 mRNA | 0.5 $\times d$ | t^-1^ |
| $k_{3dt}$ | Degradation rate of Cdh1 total | 1.0 $\times d$ | t^-1^ |
| $k_{5am}$ | Basal synthesis rate of Cdc20 mRNA | 0.005 $\times d$ | c t^-1^ |
| $k_{5bm}$ | CycB-dependent transcription rate of Cdc20 mRNA | 0.2 $\times d$ | c t^-1^ |
| $k_{5dm}$ | Degradation rate of Cdc20 mRNA | 1.386 $\times d$ | t^-1^ |
| $j_{5}$ | Hill coefficient for CycB dependent transcription rate of Cdc20 mRNA | 0.3 | c |
| $n$ | Hill constant for CycB dependent transcription rate of cdc20 mRNA | 4.0 | - |
| $k_{5a}$ | Synthesis rate of Cdc20 total | 1.0$\times d$ | t^-1^ |
| $k_{6}$ | Degradation rate of Cdc20 total | 0.05 $\times d$ | t^-1^ |
| $k_{7}$ | Activation rate of Cdc20A | 1.4 $\times d$ | t^-1^ |
| $k_{8}$ | Degradation rate of Cdc20A | 0.5$\times d$ | c t^-1^ |
| $j_{7}$ | Michaelis Menten equation for Activation rate of Cdc20A | 0.001 | c |
| $j_{8}$ | Michaelis Menten equation for Activation rate of Cdc20A | 0.001 | c |
| $k_{9}$ | Synthesis rate of IEP | 0.1$\times d$ | (c t)^-1^ |
| $k_{10}$ | Degradation rate of IEP | 0.02 $\times d$ | t^-1^ |
| $k_{11}$ | Synthesis rate of X | 0.045$\times d$ | c t^-1^ |
| $k_{12}$ | CycB mediated degradation rate of X | 2.27 $\times d$ | (c t)^-1^ |
| $k_{13}$ | Degradation rate of X | 0.004$\times d$ | t^-1^ |
| $k_{3}$ | Basal synthesis rate of Cdh1 A | 1.28$\times d$ | c t^-1^ |
| $GF$ | Growth factor | 2.0 |  |
| $k_{mm}$ | Michalis Menten constant for Growth factor-induced transcription of CycB mRNA | 0.2 |  |
| $k_{5cm}$ | Michalis Menten constant for Growth factor inhibited transcription of Cdc20 mRNA | 1.0 |  |
| $j_{5c}$ | Effective GF concentration for inhibition of Cdc20 mRNA | 0.02 |  |
| $d$ | Scaling factor for cell cycle period adjustment | 2.8 |  |

For Fig.3 d value changed – T_cc_ = 16h (d=4.35) , T_cc_ = 20h (d=3.5), T_cc_ = 24h (d=2.8), T_cc_ = 28h (d=2.52)

For Fig.4 – (A) 16h (9h+7h) – $k_{4}= 15 \left( t^{-1} \right), k_{1}=1.1{(t}^{-1})$

(B) 16h (3h+13h) -$k_{4}= 45 \left( t^{-1} \right), k_{1}=0.45{(t}^{-1}) k_{2b}= 1.8$
(C) 28hr (16h+12h) – $k_{4}= 20 \left( t^{-1} \right), k_{1}=0.6{(t}^{-1})$

(D) 28 hr (9h+19h) - $k_{4}=45 \left( t^{-1} \right), k_{1}=0.43{(t}^{-1}), k_{2b}=1.8 \left( c t \right)^{-1} d=1.8$

Only the parameters given here are changed, other parameters are kept the same as in Table 2.

**Table 3 - Abbreviated Name and description of variables.**

| **Abbreviated form** | **Description** | **Initial condition (in Concentration unit)** |
| --- | --- | --- |
| $CycB_{m}$ | CycB mRNA | 0.05 |
| $CycB$ | CycB protein | 0.3 |
| $C{20}_{m}$ | Cdc20 mRNA | 0.07 |
| $CDC20t$ | Cdc20 protein total | 0.59 |
| $Cdh_{m}$ | Cdh1 mRNA | 0.99 |
| $Cdht$ | Cdh1 protein total | 0.99 |
| $CdhA$ | The active form of Cdh1 | 0.006 |
| $Cdc20A$ | Active form of Cdc20 | 0.036 |
| $IEP$ | Intermediary Enzyme | 0.41 |

**Model Description :**

In our minimalistic mathematical model adapted from Tyson et al. 2001, we have modified the original network by incorporating the mRNA of each protein and making it growth factor (GF) dependent. The model has three regulatory components of the cell cycle CycB, Cdh1, and Cdc20. In the cell cycle, Cdh1/APC is expressed during the G1 phase, while CycB/CDKs, the activator of the cell cycle, remain high during the S, G2, and M phases of the cell cycle. We have translated all the molecular interactions depicted in **SFig 1** into ordinary differential equations (**Table 1**). **Eq. 1** shows the dynamics of CycB and cell cycle initiation by growth factor. Here we have included the correlated fluctuations in the transcription rate of CycB mRNA (1^st^ term in **Eq.1**) and its normal degradation (2^nd^ term in **Eq.1**).

$\frac{dCycBm}{dt}=\frac{\left( k_{1m}+\varepsilon_{ou1} \right)\times GF}{K_{mm}+k_{eff}*GF}-k_{1dm}\times CycBm$	**Eq. 1**

**Eq. 2** shows the dynamics of translation of CycB protein (1^st^ term in **Eq.2**), natural degradation (2^nd^ term in **Eq.1**), and Cdh1 mediated degradation of CycB (3^rd^ term in **Eq.2**).

$\frac{dCycB}{dt}=k_{1}\times CycBm-k_{2a}\times CycB-k_{2b}\times CycB\times Cdh1$	**Eq. 2**

**Eq. 3** captures the dynamics of Cdc20 mRNA; transcription is turned on by CycB independent (1^st^ term in **Eq.3**) and CycB dependent (2^nd^ term in **Eq.3**) with Hill function parameterized by J_5_ and n. Degradation of Cdc20 mRNA shown in last term of **Eq.3**.

$\frac{dCdc20m}{dt}=\left( k_{5am}+\varepsilon_{ou2} \right)+\frac{\left( k_{5bm}\boldsymbol{+}\varepsilon_{ou3} \right)}{k_{5cm}+GF \times J_{5c}}\times\frac{CycB^{n}}{J_{5}^{n}+CycB^{n}}-k_{5dm}\times Cdc20m$	**Eq. 3**

**Eq.4** and **Eq.5** show the total Cdc20 protein and Cdc20A dynamics, respectively. The concentration of total Cdc20, including both active and inactive forms, has translation and degradation terms **Eq.4**). Newly synthesized Cdc20 is initially inactive (Cdc20 total – Cdc20 active), then activated by IEP (Shown in **Eq. 9**) with some intermediate steps (1st term in **Eq.5**). Cdc20 activation ensures the DNA synthesis and chromosome alignment before anaphase, if they are not finished on time, then spindle checkpoint protein MAD will inactivate the Cdc20 (2^nd^ term in **Eq.5**)

$\frac{dCdc20T}{dt}=k_{5a}\times Cdc20m-k_{6}\times Cdc20T$	**Eq. 4**

$$\frac{dCdc20A}{dt}=k_{7}\times IEP\times\frac{Cdc20T-Cdc20A}{J_{7}+Cdc20T - Cdc20A}-k_{8}\times Mad\times\frac{Cdc20A}{J_{8}+Cdc20A} -k_{6}\times Cdc20A$$

**Eq. 5**

We have modeled the Cdh1 mRNA in **Eq. 6**, we incorporated the colored noise in the transcription rate (1^st^ term in **Eq.6**). Last term shows the natural degradation of Cdh mRNA.

$\frac{dCdhm}{dt}=\left( k_{3m}+\varepsilon_{ou4} \right)-k_{3dm}\times Cdhm$	**Eq.6**

**Eq. 7** and **Eq.8** capture the total Cdh1 protein and active Cdh1 protein dynamics, respectively. **Eq. 7** includes a translation and a degradation term for total Cdh1 protein. **Eq. 8** has an activation and deactivation mechanism of Cdh1, which is modeled using Michaelis-Menten kinetics. The activation of Cdh1 is driven by Cdc20A (active form) (1^st^ term in **Eq.8**). meanwhile, CycB deactivates the active form of Cdh1 through phosphorylation (2^nd^ term in **Eq. 8**).

${\frac{dCdht}{dt}=k}_{3a}\times Cdhm-k_{3dt}\times Cdht$	**Eq.7**

$$\frac{dCdhA}{dt}=\left( k_{3}+k_{3b}\times Cdc20A \right)\times\frac{Cdht-Cdh1}{J_{3}+Cdht-Cdh1}-k_{4}\times CycB\times\frac{Cdh1}{J_{4}+Cdh1}-k_{3dt}\times Cdh1$$

**Eq.8**

**Eq.9** includes the dynamics of the hypothetical intermediary enzyme (IEP); IEP generates a time lag between CycB activation and Cdc20 rise. The first term shows the activation of IEP by CycB where total concentration is scaled to 1. The last term depicts the inactivation of IEP.

$\frac{dIEP}{dt}=k_{9}\times CycB\times(1-IEP)-k_{10}\times IEP$	**Eq.9**

The structure of dynamical equations (**Eq.1 - Eq.9**) remains consistent throughout our simulations. Parameter values for the ODEs were directly taken from Tyson et al. 2001, and parameters related to mRNAs were chosen to match the protein time profiles from Tyson et al. (**Table 2**). To vary the cell cycle durations and cell cycle phase durations, we adjusted a few parameters from those listed in Table 2; these changes are provided in the notes below Table 2.

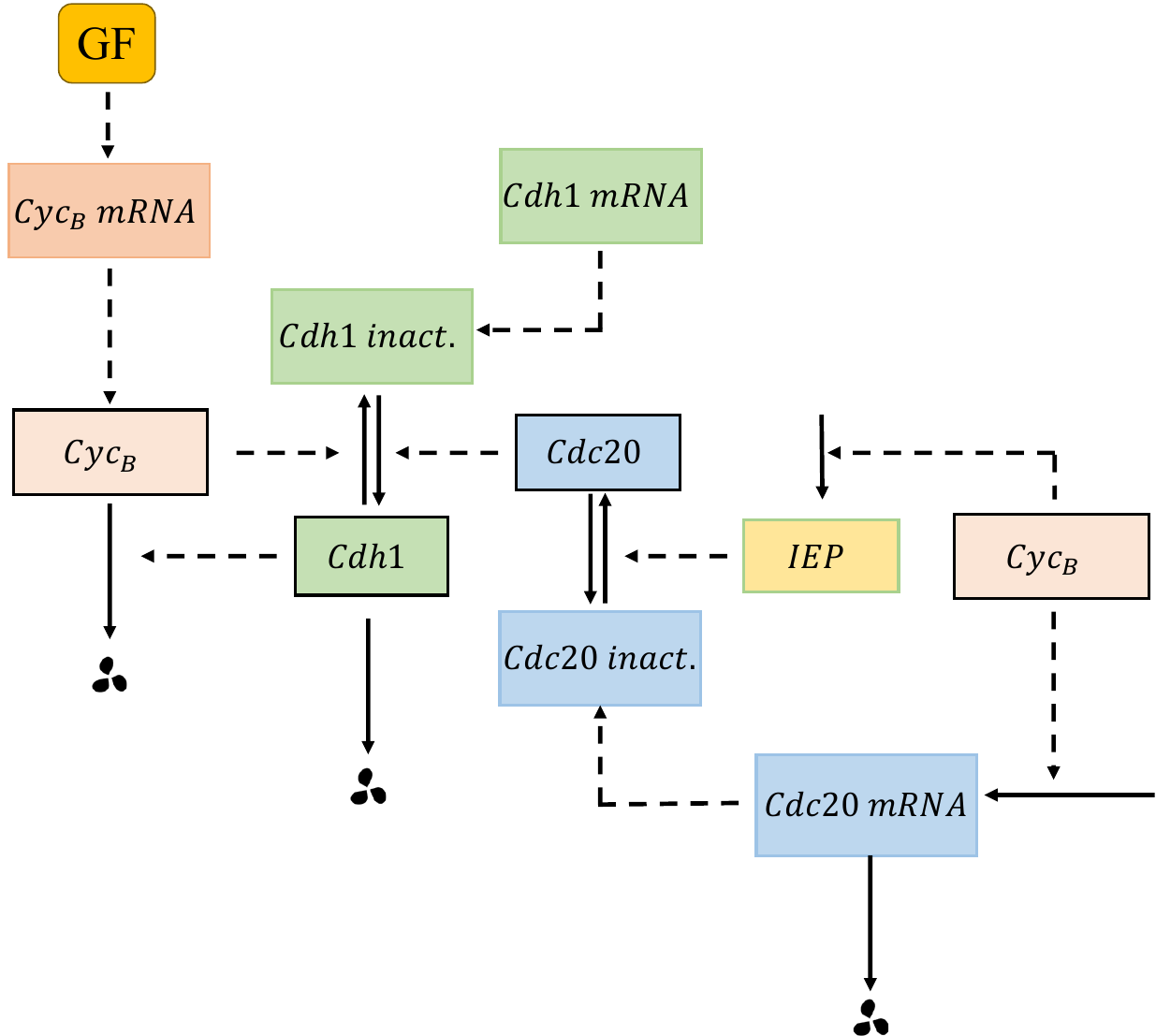

**SFig.1** Detailed cell cycle regulatory network (adapted from Tyson et al. 2001).

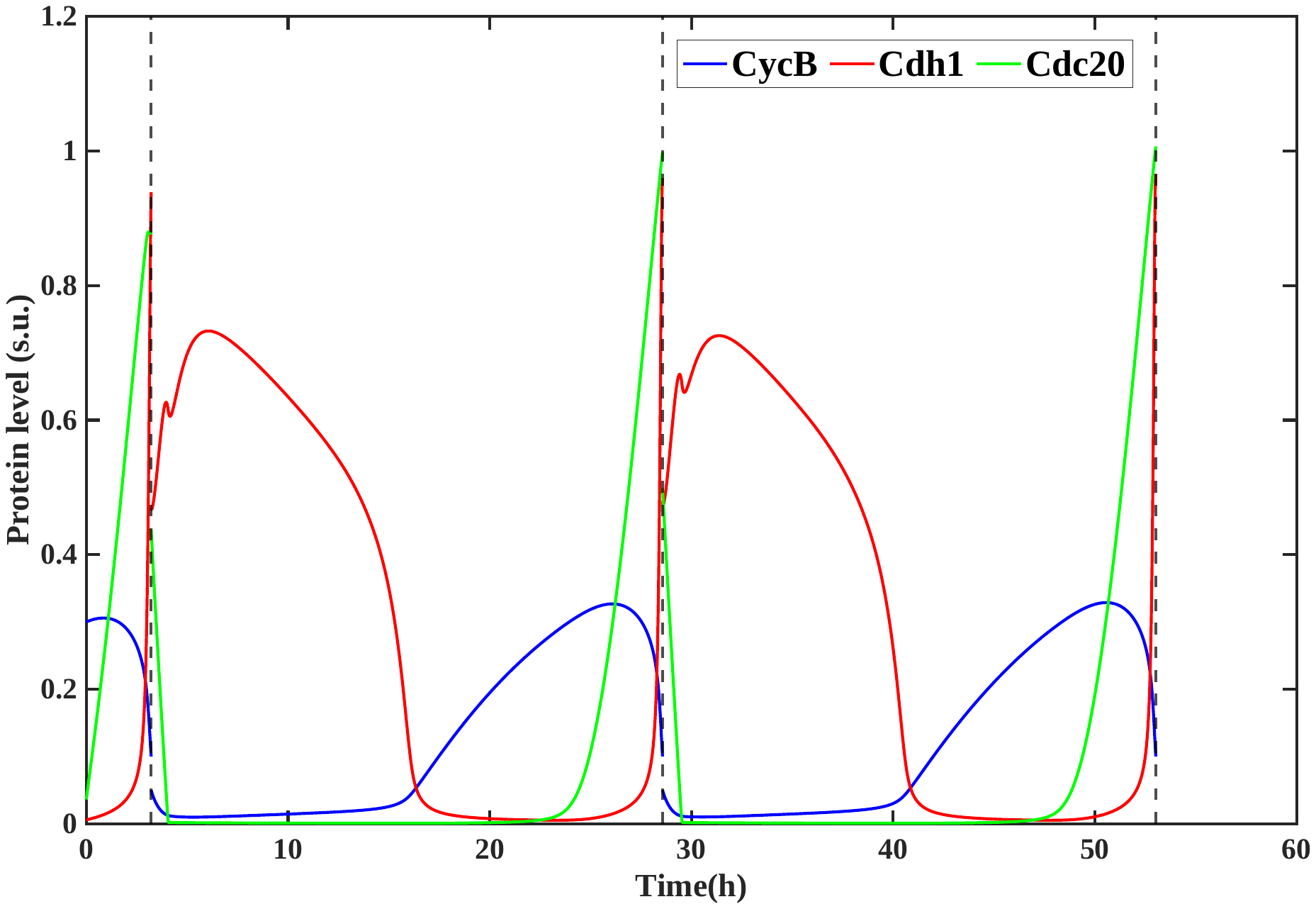

**SFig.2** Temporal dynamics of CycB, Cdh1 and Cdc20 Proteins. (Vertical dotted line shows the cell division event)

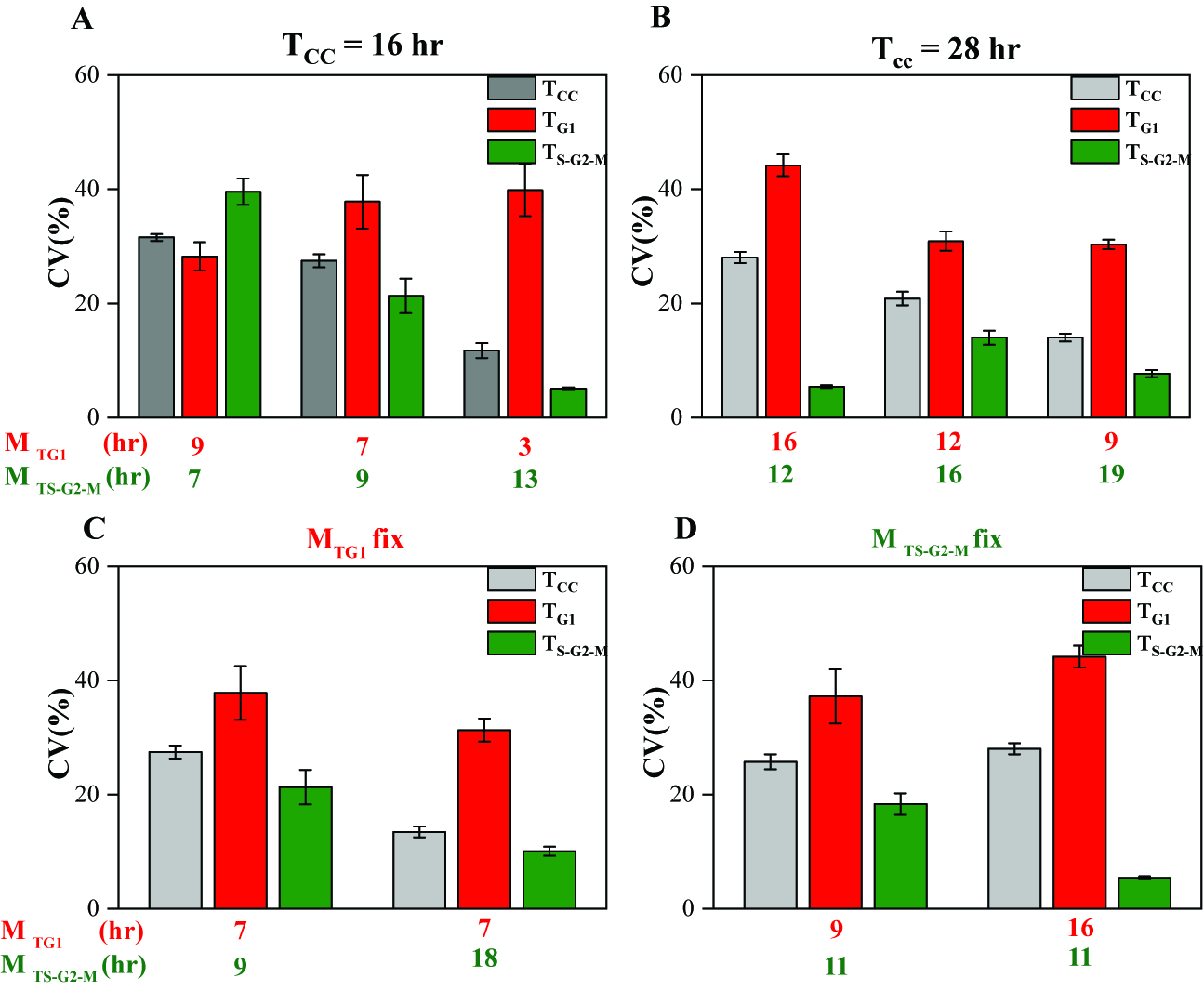

**SFig.3** Average variability (%CVs) of T_CC_, T_G1,_ and T_S-G2-M_ distributions at (A) 16 hr cell cycle duration with three combinations of mean G1 and S-G2-M durations (M_TG1_ = 9hr + M_TS-G2-M_ = 7hr (left), M_TG1_=7hr + M_TS-G2-M_ = 9hr (center), M_TG1_ = 3hr + M_TS-G2-M_ = 13 hr (right)). (B) 28 hr cell cycle duration with three combinations of mean G1 and S-G2-M durations (M_TG1_ = 16hr + M_TS-G2-M_ = 12hr (left), M_TG1_=12hr + M_TS-G2-M_ = 16hr (center), M_TG1_ = 9hr + M_TS-G2-M_ = 19 hr (right)). (C) 7 hr M_TG1_ with 9hr (left) and 18hr (right) M_TS-G2-M._ (D) 11hr M_TS-G2-M_ with 9hr (left) and 16hr (right) M_TG1_ duration. (Mean CV calculated from three replicates each containing 300 cells, error bar represents the standard deviation)

**Significance Analysis** - We conducted a significance analysis to evaluate whether the observed correlations between specific lineage pairs were significantly different. Specifically, we compared the correlation coefficients (R) among various types of pairs— (i) D-D M-D, (ii) D-D C-C, (iii) M-D C-C (**SFig.4-7**) — across three individual replicates, each containing 300 cells.

For each cell subpopulation in each replicate, we generated 500 bootstrap random samples and calculated the correlation coefficient (R). To assess the statistical significance, we applied Fisher's z-transformation to the Pearson correlation coefficient values and then calculated p-values using the Student's t-test.

The figures (**SFig.4-7**) show p-value plots, where p-values less than 0.05 indicate statistically significant differences in the correlation coefficients of the lineage pairs, conversely p-values greater than 0.05 suggest that the difference in correlation coefficients of lineage pairs are not statistically significant.

Our significant analysis shows that D-D correlations significantly differ from M-D and C-C correlations (p<0.05). Whereas in most cases, C-C and M-D pairs also show considerable differences (p<0.05) in both scenarios where CMI is satisfied, and CMI is not satisfied.

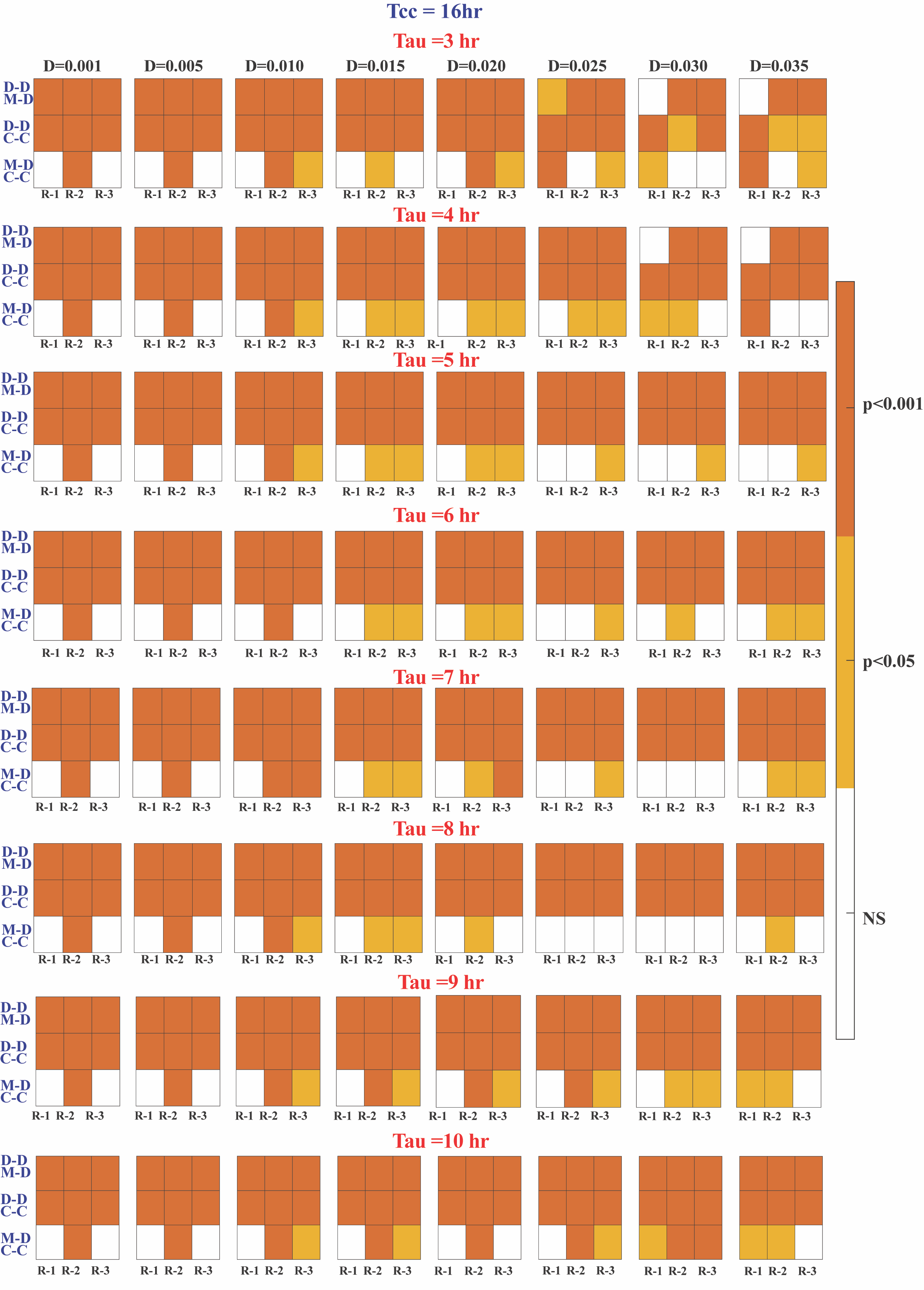

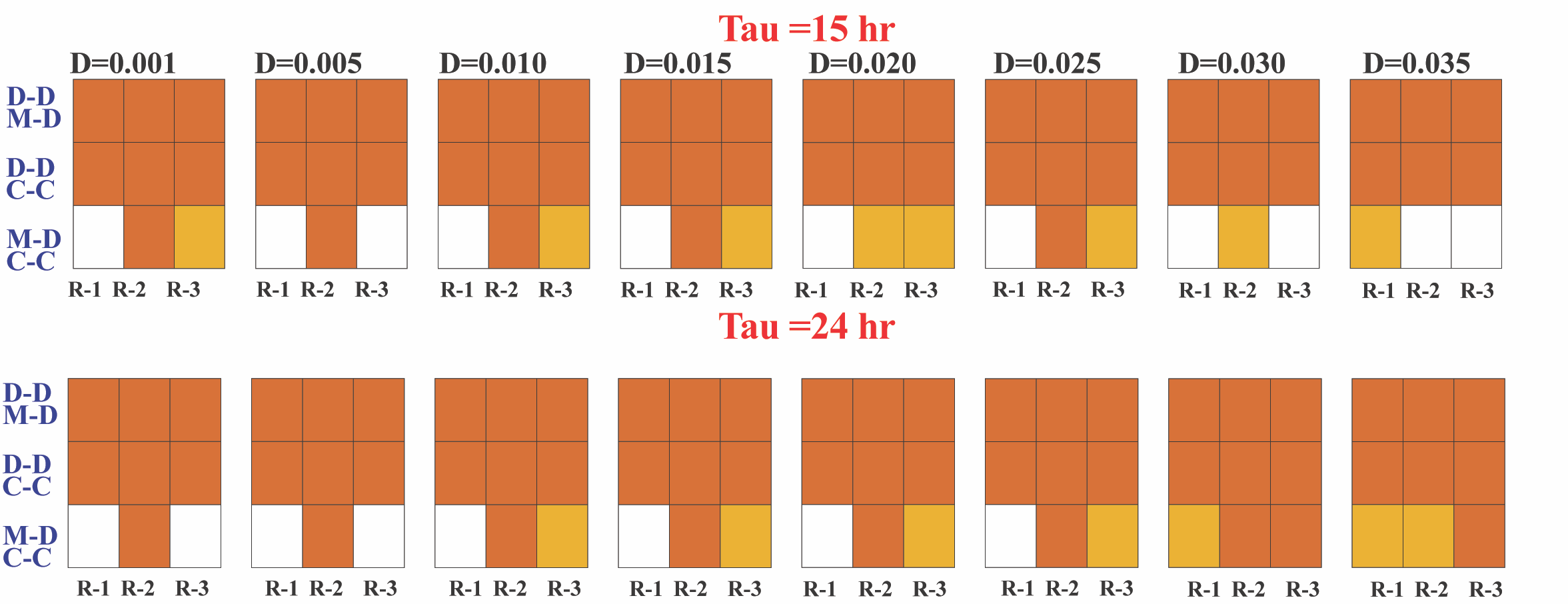

**SFig.4** Significance analysis (For **Fig 3A(i) & Fig 3B(i)**) of correlations between lineage pairs of individual replicates for different tau values 3hr to 10hr, 15hr and 24 hr. (Colour bar in right shows- **NS** (Not Significant), **p<0.05** and **p<0.001**). The total cell cycle time is **16 hr** here.

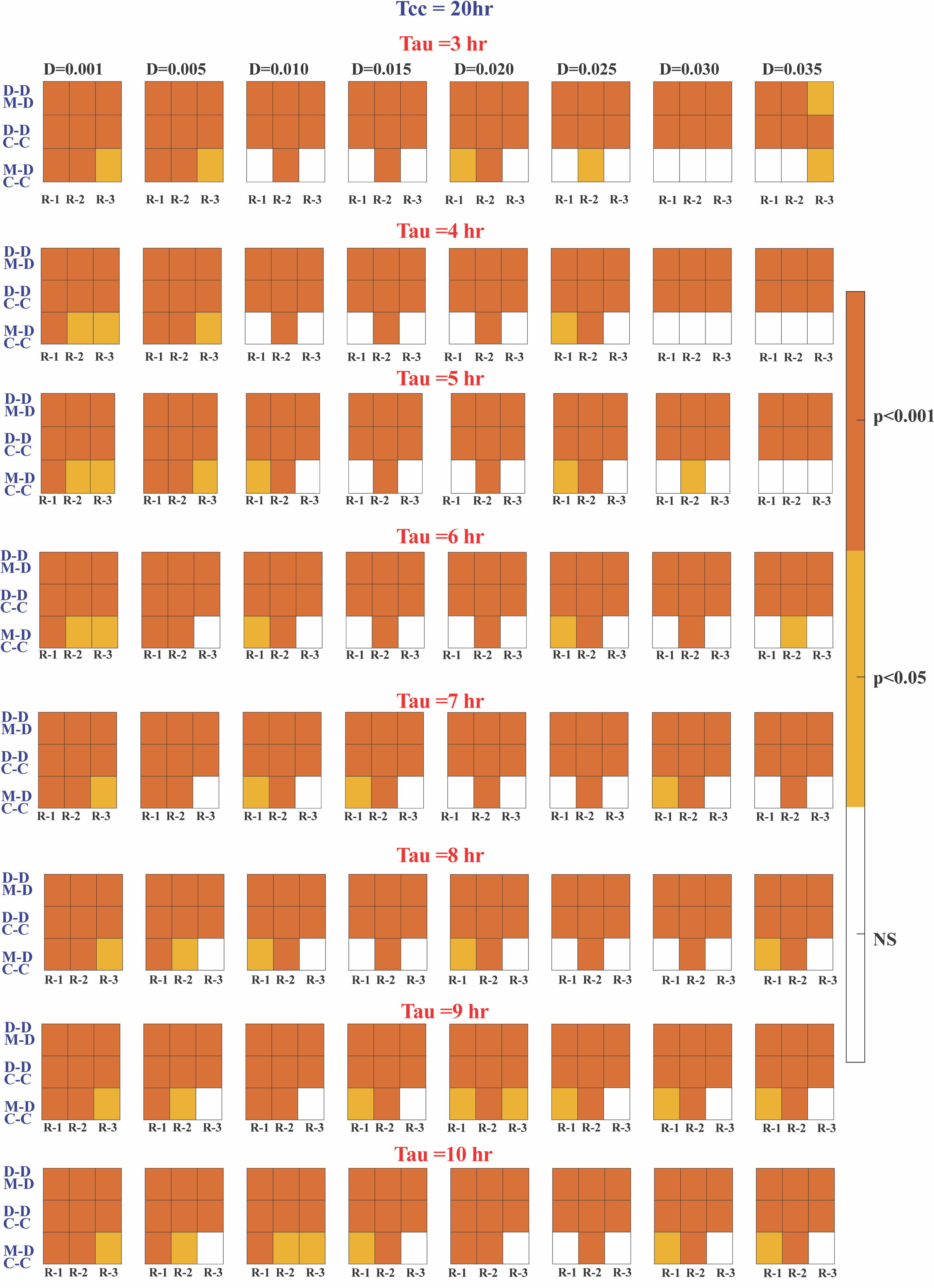

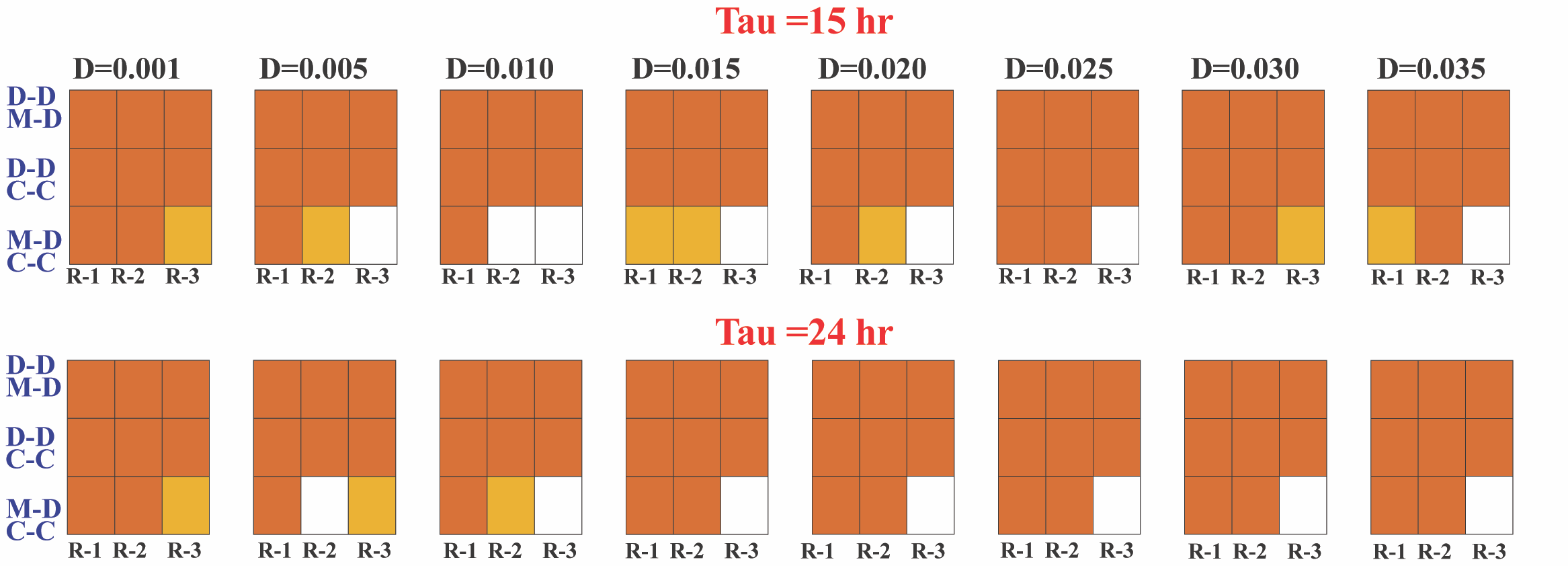

**SFig.5** Significance analysis (For **Fig 3A(ii) & 3B(ii)**) of correlations between lineage pairs of individual replicates for different tau values 3hr to 10hr, 15hr and 24 hr. (Colour bar in right shows- **NS** (Not Significant), **p<0.05** and **p<0.001**). The total cell cycle time is **20 hr** here.

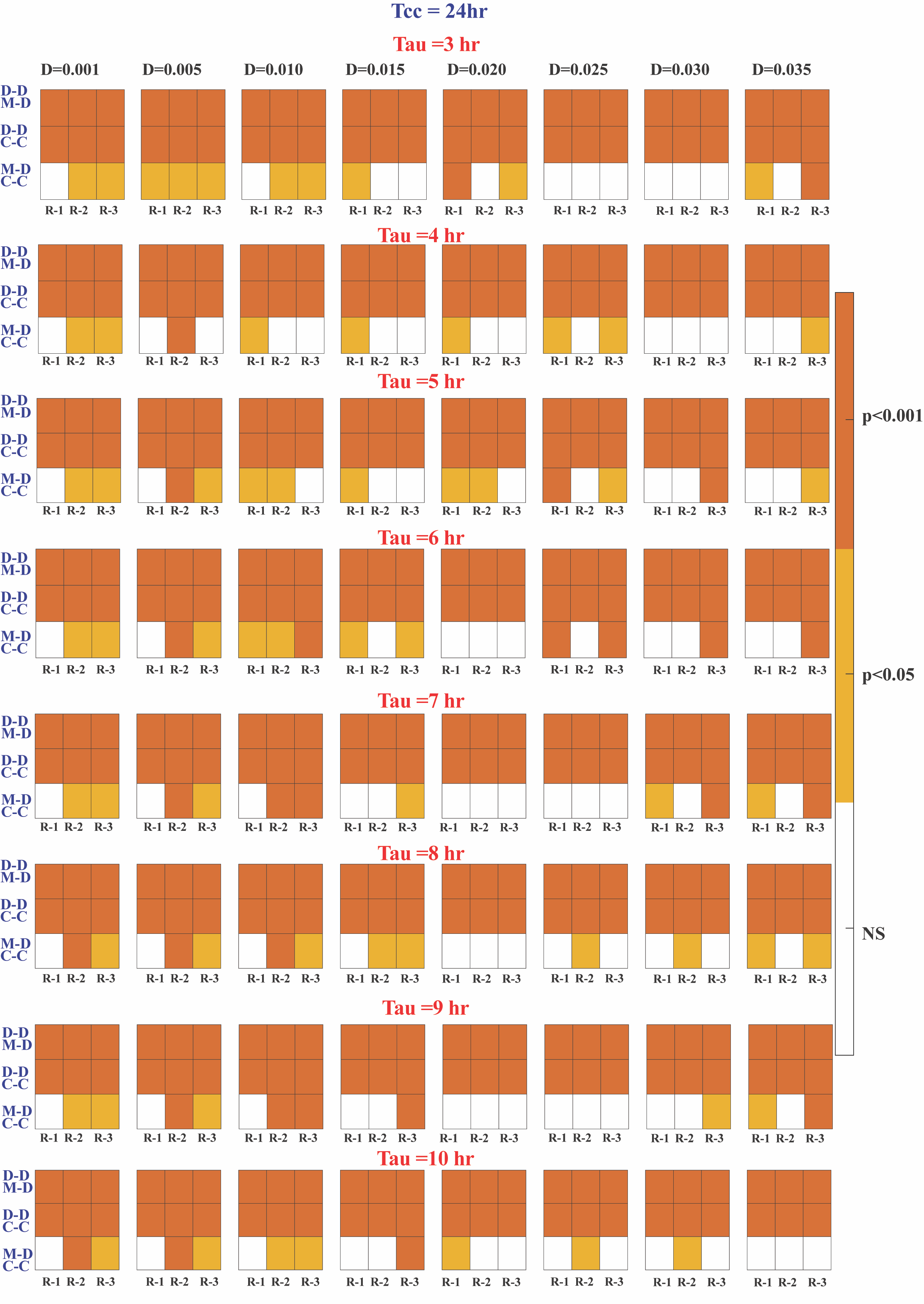

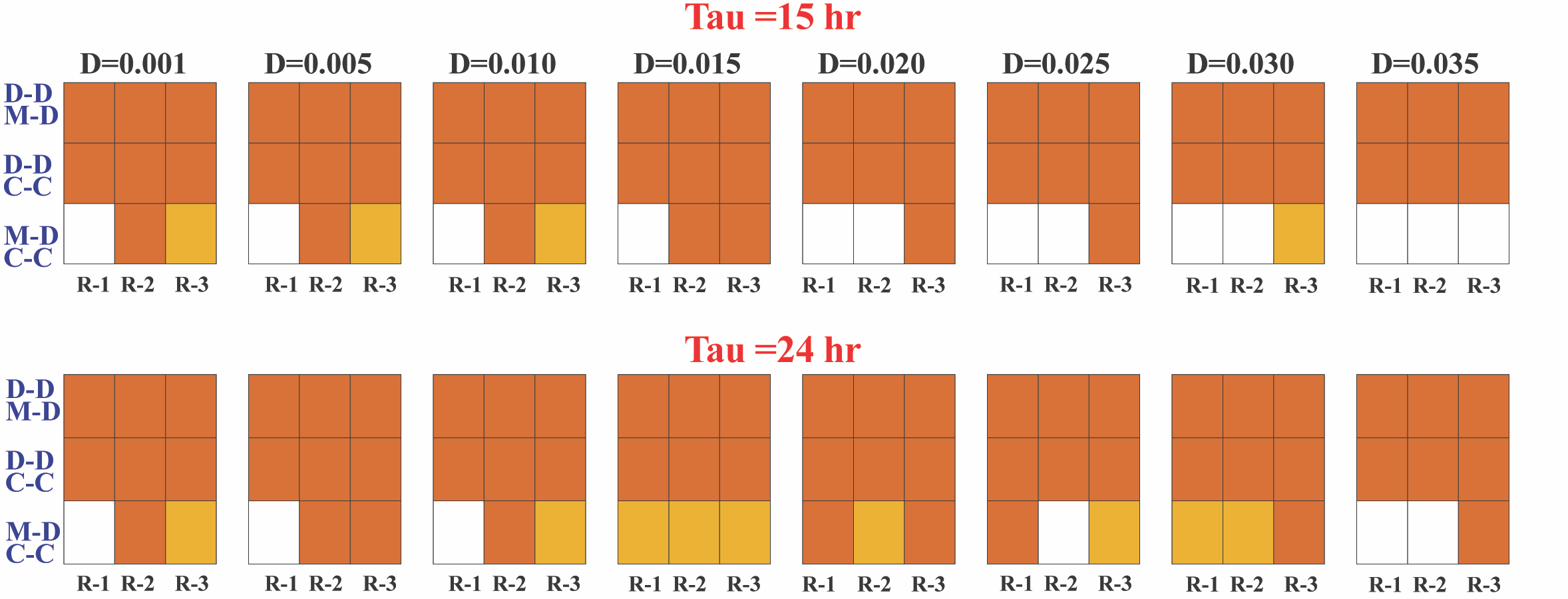

**SFig.6** Significance analysis (For **Fig 2D(i),** **Fig 3A(iii) & Fig 3B(iii)**) of correlations between lineage pairs of individual replicates for different tau values 3hr to 10hr, 15hr and 24 hr. (Colour bar in right shows- **NS** (Not Significant), **p<0.05** and **p<0.001**). The total cell cycle time is **24 hr** here.

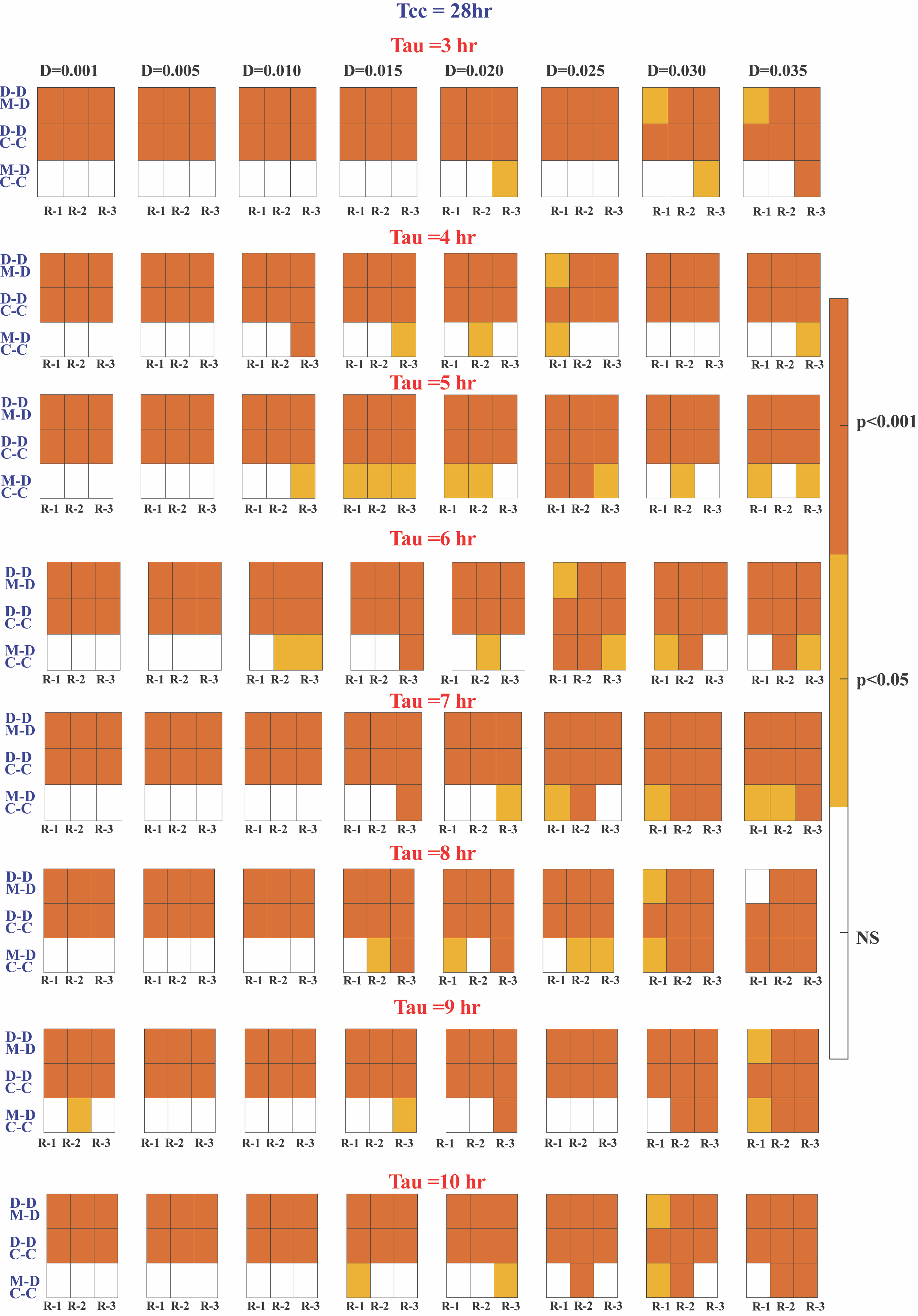

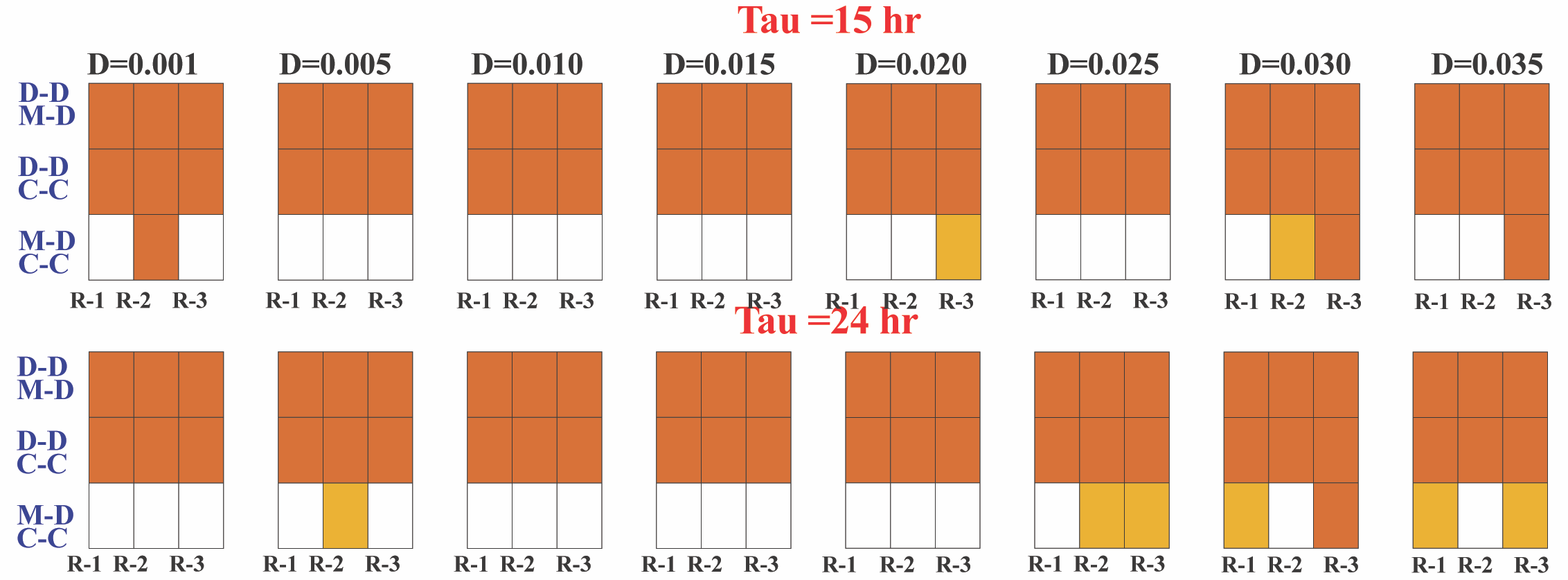

**SFig.7** Significance analysis (For **Fig 3A(iv) & Fig 3B(iv)**) of correlations between lineage pairs of individual replicates for different tau values 3hr to 10hr, 15hr and 24 hr. (Colour bar in right shows-**NS** (Not Significant), **p<0.05** and **p<0.001**). The total cell cycle duration is **28 hr** here.
